# Hepatic Vagal Afferents Convey Clock-Dependent Signals to Regulate Circadian Food Intake

**DOI:** 10.1101/2023.11.30.568080

**Authors:** Lauren N. Woodie, Lily C. Melink, Mohit Midha, Alan M. de Araújo, Caroline E. Geisler, Ahren J. Alberto, Brianna M. Krusen, Delaine M. Zundell, Guillaume de Lartigue, Matthew R. Hayes, Mitchell A. Lazar

## Abstract

Circadian desynchrony induced by shiftwork or jetlag is detrimental to metabolic health, but how synchronous/desynchronous signals are transmitted among tissues is unknown. Here we report that liver molecular clock dysfunction is signaled to the brain via the hepatic vagal afferent nerve (HVAN), leading to altered food intake patterns that are corrected by ablation of the HVAN. Hepatic branch vagotomy also prevents food intake disruptions induced by high-fat diet feeding and reduces body weight gain. Our findings reveal a previously unrecognized homeostatic feedback signal that relies on synchrony between the liver and the brain to control circadian food intake patterns. This identifies the hepatic vagus nerve as a therapeutic target for obesity in the setting of chrono-disruption.

**One Sentence Summary:** The hepatic vagal afferent nerve signals internal circadian desynchrony between the brain and liver to induce maladaptive food intake patterns.

## Main Text

Mammalian circadian rhythms are arranged in a hierarchical order with the suprachiasmatic nucleus (SCN) functioning as the master circadian clock. Light cues set SCN rhythms to a ∼24-hour cycle by initiating a transcription/translation feedback loop (TTFL) of molecular clock genes comprised of transcriptional activators (e.g. BMAL1) and transcriptional repressors (e.g. REV-ERBs) (*1*). Although the SCN is regarded as the master clock, almost every cell in the body possesses an oscillating TTFL (*1*). Indeed, the liver TTFL is highly sensitive to fed/fasting rhythms and is an established food-entrainable oscillator (*2–5*).

Synchrony between the light-entrained SCN and food-entrained liver is critical for maintaining chrono-metabolic homeostasis in the face of varying environmental cues (*6*– *9*). Rodent and human studies have shown that desynchrony between these tissues is detrimental to organismal health (*6*, *7*, *10*). Although the consequences of circadian desynchrony are well known, the mechanism through which synchrony and desynchrony are communicated within an organism remains a major unanswered question.

The REV-ERBα/β nuclear receptors are important regulatory components of chrono-metabolic homeostasis and their deletion induces desynchrony within an organism (*3*, *11–14*). Thus, to investigate the ways through which the liver and brain communicate circadian signals we induced internal desynchrony by deleting REV-ERBα and REV-ERBβ in hepatocytes (hepatocyte REV-ERB double knockout; HepDKO) by performing tail vein injections of AAV8-TBG-Cre on adult REV-ERBα/β floxed mice (**Fig. 1A**). This model eliminates potential developmental effects of the knockout and allows for the interrogation of internal desynchrony, specifically between the brain and liver, without confounding effects of manipulating the clock in other tissues (*3*). The peak of REV-ERBs expression in the mouse liver is Zeitgeber time (ZT) 10 and *Nr1d1* and *Nr1d2* (REV-ERBα and REV-ERBβ, respectively) expression in the livers of HepDKO mice were nearly undetectable at that time (**Fig. 1B-C**) (*3*). As expected, loss of hepatocyte REV-ERBs led to marked derepression of the positive clock output gene *Arntl* (BMAL1) at ZT10, when physiological expression of REV-ERBs is maximal (**Fig. 1D**)

**Figure 1.**
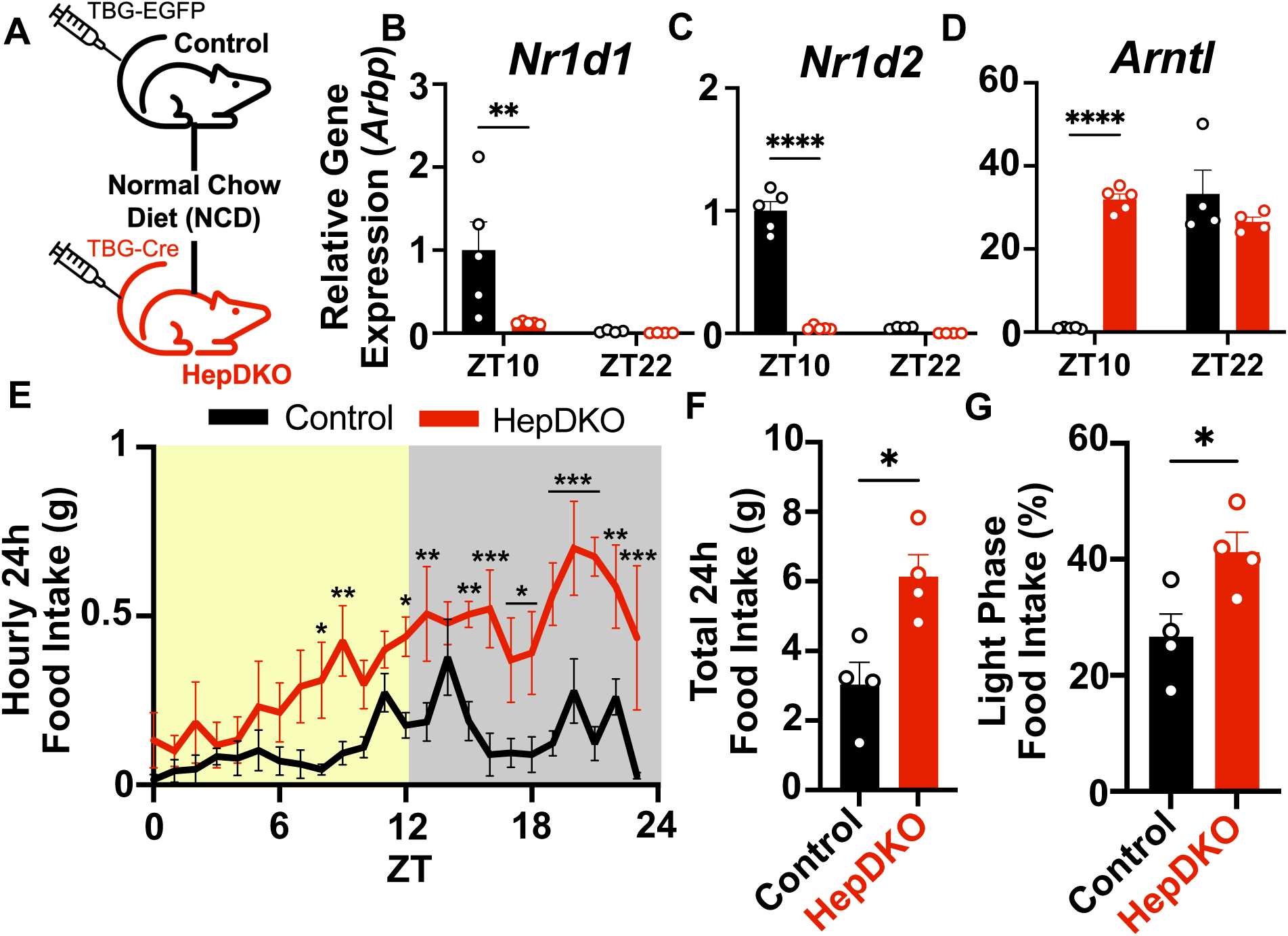
Internal desynchrony induced by loss of hepatocyte REV-ERBs (HepDKO) disrupts food intake patterns. (**A**) Hepatocyte specific deletion of REV-ERBs was achieved by i.v. injection of male *Nr1d1^fl/lf^/Nr1d2^fl/fl^* mice with AAV8-TBG-Cre. The control group was created by i.v. injection of *Nr1d1^fl/fl^/Nr1d2^fl/fl^* littermates with AAV8-TBG-EGFP. (**B-D**) Confirmation of REV-ERBs deletion in the liver and its effect on BMAL1 (*Arntl*) by RT-qPCR at Zeitgeber time (ZT) 10 and 22 (n = 4-5, mean ± SEM). (**E**) Food intake every hour over 24hrs measured in control and HepDKO animals in light:dark conditions (n = 4, mean ± SEM). (**F-G**) Total 24h food intake and percentage of 24h food intake in the light phase were calculated in control and HepDKO animals (n = 4, mean ± SEM). (**B-D & F-G**) Results were compared by Mann-Whitney U test. (**E**) Results were compared by repeated measures ANOVA. *p<0.05, **p<0.01, ***p<0.001, ****p<0.0001.

Remarkably, HepDKO resulted in a striking disruption of 24-hour food intake patterns (**Fig. 1E**). HepDKO mice consumed more food over a 24-hour time period (**Fig. 1F**) and, consistent with previous reports on liver-specific knockout of molecular clock components (*13*, *15*), HepDKO animals ate a higher percentage of their total food in the light/inactive phase than control mice (**Fig. 1G, Fig. S1A**). Body weight did not change significantly four weeks after the HepDKO (**Fig. S2A)**, potentially due to compensatory elevation of oxygen consumption (VO_2_) (**Fig. S2B**). Furthermore, these changes in food intake patterns were not accompanied by altered activity over 24 hours (**Fig. S2C**).

We next considered how desynchrony between the brain and liver could confer changes in food intake patterns. The execution of food intake is a centrally regulated behavior with major centers, such as the arcuate nucleus (Arc), located in the hypothalamus (*16*). Indeed, neurons within the Arc have recently been shown to integrate past circadian food intake experience with current metabolic needs to influence feeding behaviors (*17*). Therefore, we hypothesized that circadian disruption in the HepDKO liver alters this function of the Arc to induce aberrant food intake patterns (**Fig. 2A**). Indeed, bulk RNAseq on Arc punches collected every six hours over 24-hours revealed that approximately 75% of the rhythmic Arc transcriptome was disrupted by HepDKO (**Fig. 2B-C**), although the rhythmic expression of classic molecular clock genes in the Arc was retained (**Fig. 2D**). KEGG pathway analysis revealed that disrupted genes were enriched for p53 and mTOR signaling pathways both of which have been implicated in Arc regulation of appetite and food intake (**Fig. S3A**) (*18*, *19*).

**Figure 2.**
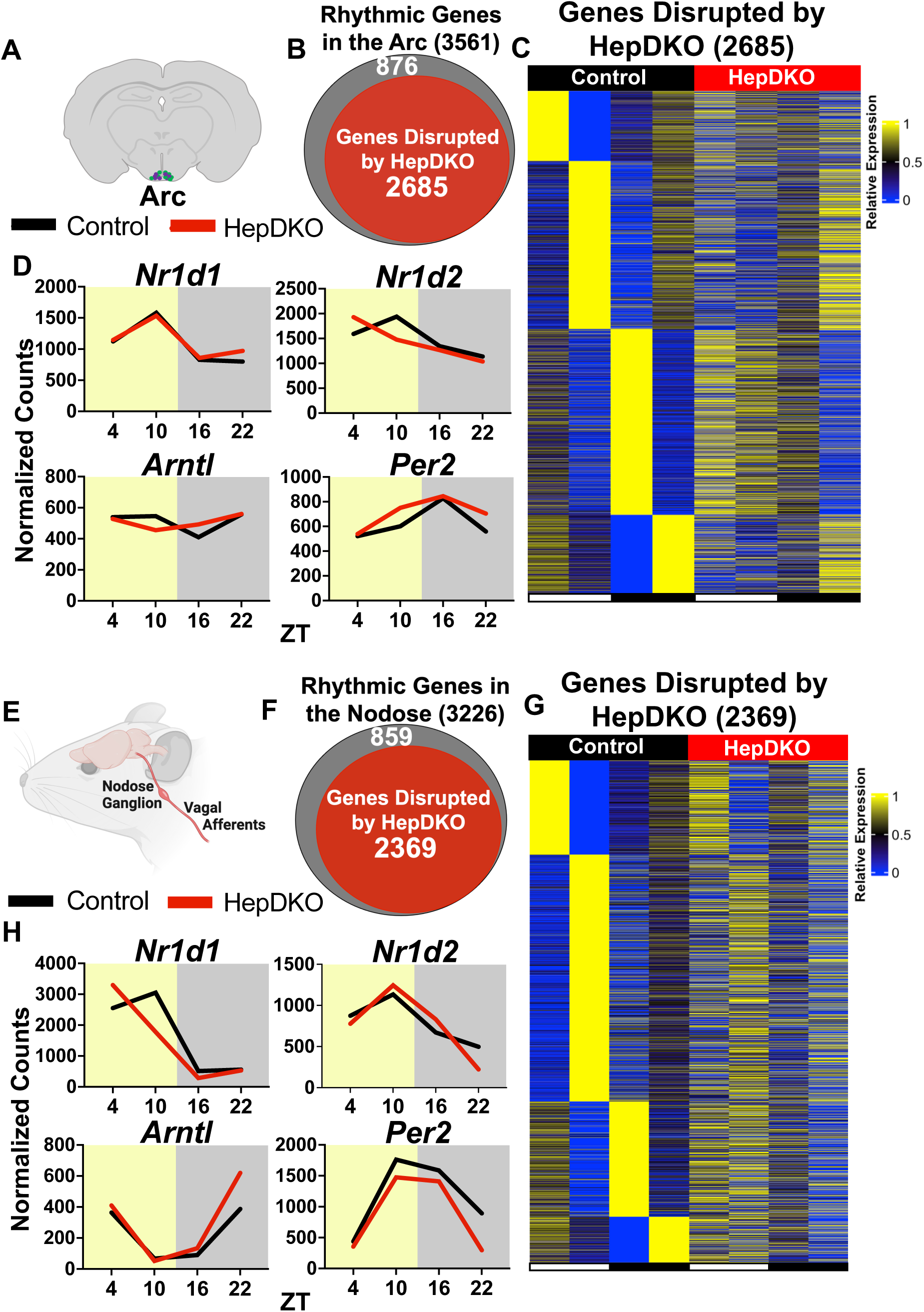
HepDKO alters the transcriptome of the arcuate nucleus (Arc) and nodose ganglia. (**A**) Site of brain micropunch location to collect Arc samples from control and HepDKO animals. (**B**) Rhythm disrupted genes identified from bulk RNAseq in the Arc of control and HepDKO animals. (**C**) Heatmap representing rhythm disrupted genes in the Arc of HepDKO animals compared to control. (**D**) The molecular clock of the Arc in control and HepDKO animals (n = 1 pool of 3 Arc per group per time point). (**E**) Site of tissue collection to harvest nodose ganglia samples from control and HepDKO animals. (**F**) Rhythm disrupted genes identified from bulk RNAseq in the nodose ganglia of control and HepDKO animals. (**G**) Heatmap representing rhythm disrupted genes in the nodose ganglia of HepDKO animals compared to control. (**H**) The molecular clock of the nodose in control and HepDKO animals (n = 1 pool of 4 left nodose per group per time point).

The Arc consists of two major cell types: agouti-related peptide (AgRP)/neuropeptide Y (NPY) releasing neurons and pre-opiomelanocortin (POMC)/ cocaine-amphetamine related transcript (CART) releasing neurons (*20*). However, circulating levels of physiological regulators of these neurons including glucose, insulin, and leptin were not significantly changed in HepDKO mice at ZT10 (**Fig. S4A-C**). Therefore, we considered whether HepDKO altered Arc activity via a neural signal.

The liver has been shown to relay metabolic signals to the brain via the hepatic vagus nerve (HV) (*21–26*) and vagal signaling has been implicated in controlling allostatic behaviors in response to varying metabolic states (*27*). Thus, we hypothesized that the HV may relay rhythmic signals from the liver to inform food intake patterns, and this may be the mechanism through which HepDKO induces aberrant feeding. The cell bodies of the hepatic vagal afferent nerve are located in the nodose ganglia which are the first sites of integration for signals from the liver (**Fig. 2E**) (*28*, *29*). To determine if HepDKO altered signaling through this neural pathway we collected nodose ganglia every six hours for 24-hours from control and HepDKO animals for bulk RNAseq. HepDKO disrupted about 73% of the rhythmic nodose ganglia transcriptome (**Fig. 2F-G**). Like the Arc, the nodose ganglia molecular clock was minimally affected by HepDKO (**Fig. 2H**). Disrupted genes were enriched for several synaptic signaling pathways such as cholinergic, glutamatergic, and serotonergic pathways (**Fig. S3B**).

To investigate the role of the HV, we next performed common hepatic branch vagotomy (HVx) on control and HepDKO animals. This surgery exclusively severs vagal innervation to the liver and a small portion of the duodenum and pancreas (*28*). We performed Sham and HVx surgeries simultaneously with i.v. injection of AAV8-TBG-EGFP or AAV8-TBG-Cre to create control and HepDKO animals, respectively, within each surgical group (**Fig. 3A**). HVx had no effect on the liver or Arc molecular clock at ZT10 or ZT22 (**Fig 3B-C, Fig. S5A-B**). Remarkably, HVx prevented HepDKO-induced disruption in daily food intake patterns (**Fig. 3D**). HVx HepDKO animals consumed less total food and shifted the majority of their food intake from the light phase to the dark phase when compared to Sham HepDKO animals (**Fig. 3E-F, Fig. S6A**).

**Figure 3.**
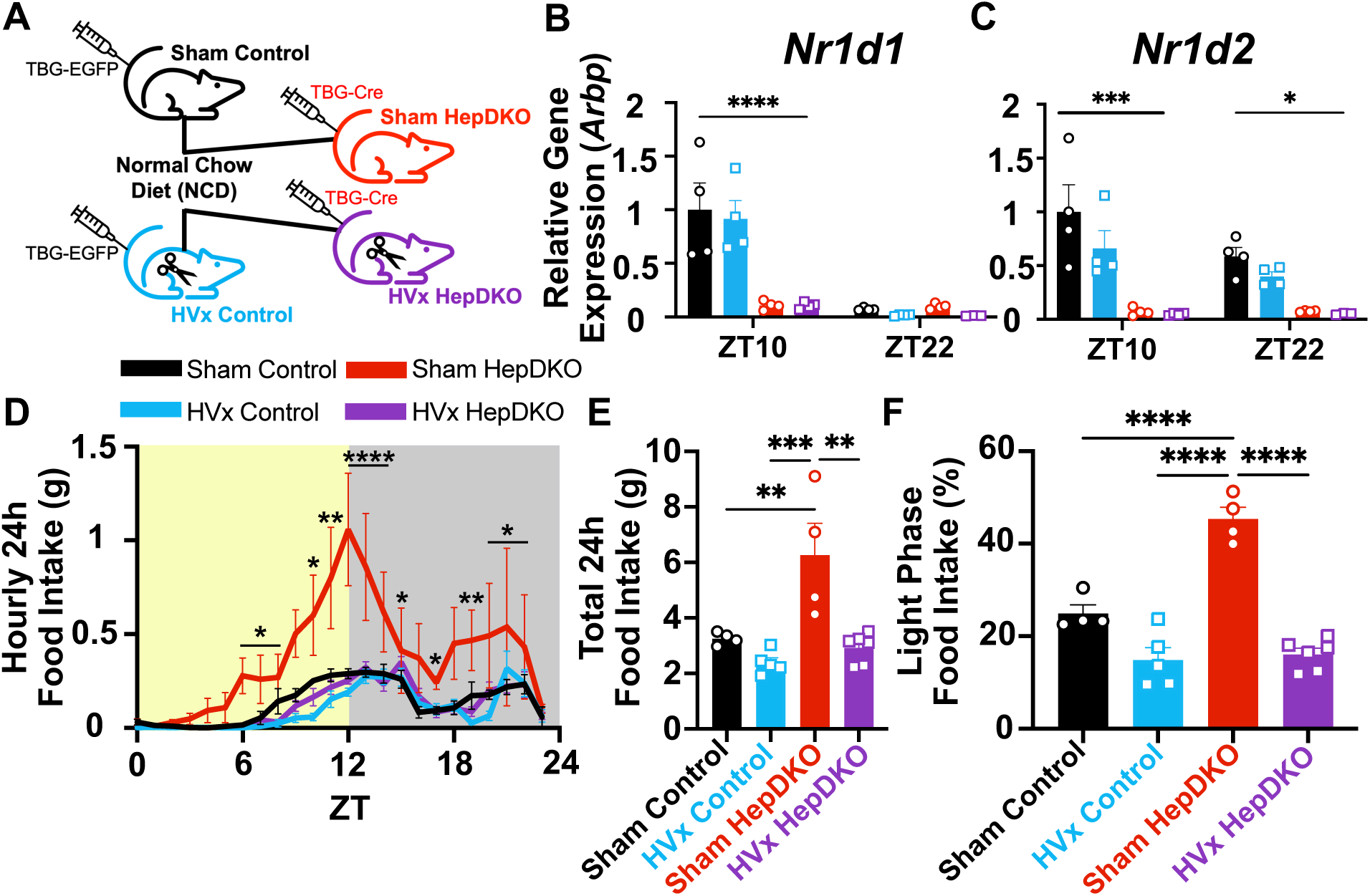
Hepatic vagotomy prevents disrupted food intake patterns from internal desynchrony induced by HepDKO. (**A**) Male *Nr1d1^fl/fl^/Nr1d2^fl/fl^*littermates received i.v. injections of either AAV8-TBG-EGFP to create control groups or AAV8-TBG-Cre to create HepDKO groups and at the same time underwent surgery to create Sham and HVx groups. (**B-C**) Confirmation of REV-ERBs (*Nr1d1* and *Nr1d2*) deletion in the liver by RT-qPCR (n = 4, mean ± SEM). (**D**) Food intake patterns every hour over 24h in light:dark conditions (n = 4-6 mean ± SEM). (**E-F**) Total 24h food intake and percentage of 24h food intake in the light phase were calculated in REV-ERB deletion and control groups (n = 4-6, mean ± SEM). (**B-C & E-F**) Results were compared by one way ANOVA. (**D**) Results were compared using a repeated measures ANOVA. *p<0.05, **p<0.01, ***p<0.001, ****p<0.0001.

We next considered whether the effect of HepDKO on food intake, and the central conveyance of this signal by the HV could be specific to the loss of REV-ERBs in hepatocytes, or an effect of general hepatic clock disruption. To address this, we deleted the positive clock output protein BMAL1 in hepatocytes of adult mice (HepBKO) by i.v. injection of AAV8-TBG-EGFP or AAV8-TBG-Cre into *Arntl^fl/fl^* animals (**Fig. 4A**). Simultaneous with viral injection, we performed Sham and HVx surgeries in the control and HepBKO animals to test the role of hepatic vagal signaling, (**Fig. 4A**). Deletion of BMAL1 was confirmed by RT-qPCR (**Fig. 4B-C, Fig. S7A**) and unchanged expression of the Arc molecular clock in all groups was confirmed by RT-qPCR (**Fig. S7B**). Like HepDKO mice, HepBKO mice exhibited disrupted daily patterns of food intake, with a trend toward increased percentage of total food take in the light/inactive period, which was similarly prevented by HVx (**Fig. 4D-F, Fig. S8A**). These data strongly suggest that disruption of the hepatocyte clock, rather than the loss of REV-ERBs per se, induces vagally mediated alterations in food intake patterns.

**Figure 4.**
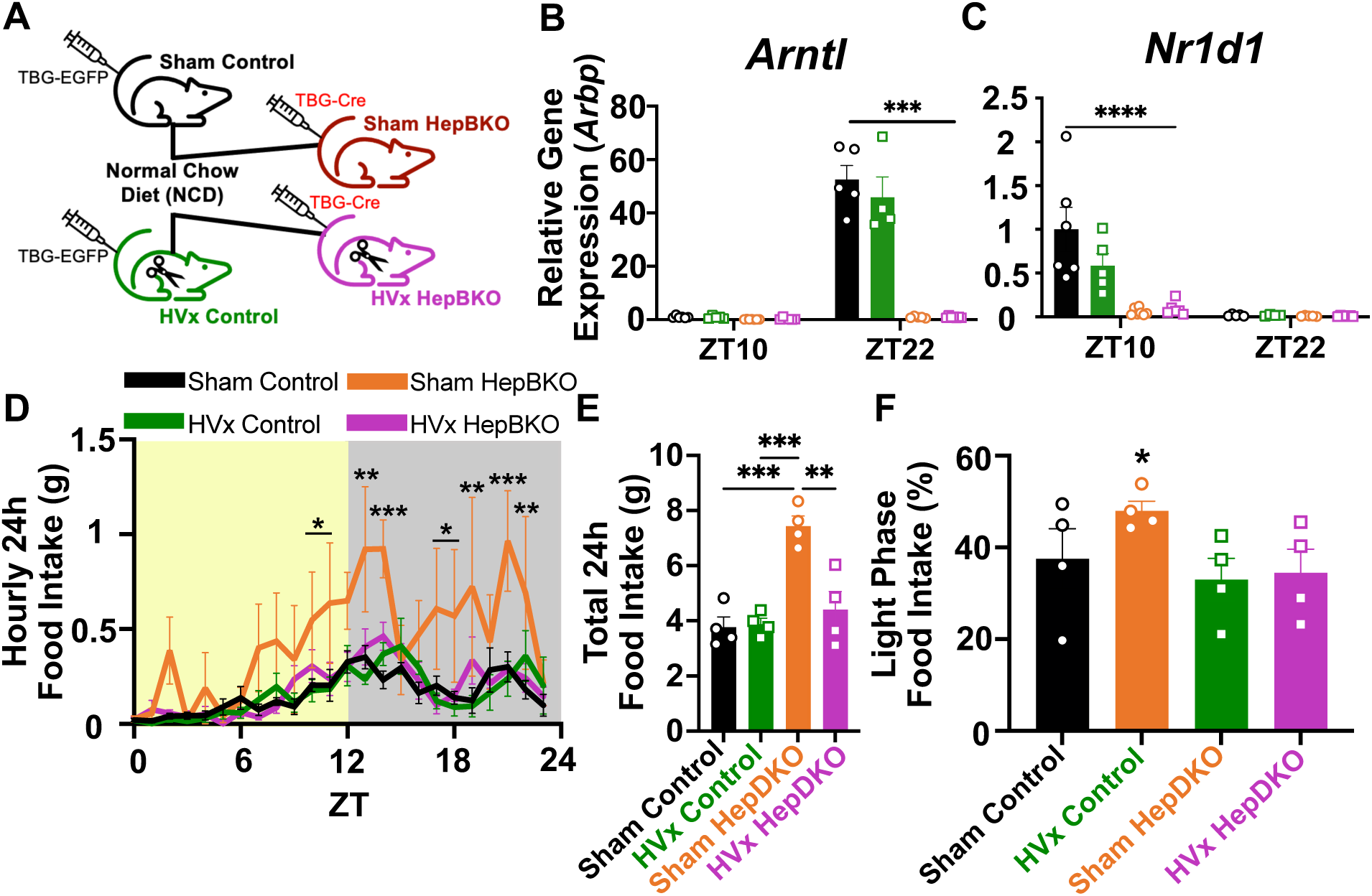
Hepatic vagotomy prevents disrupted food intake patterns from internal desynchrony induced by HepBKO. (**A**) Male *Arntl^fl/fl^* littermates received i.v. injections of either AAV8-TBG-EGFP to create control groups or AAV8-TBG-Cre to create HepBKO groups at the same time as surgery to create Sham and HVx groups. (**B-C**) Confirmation of BMAL1 (*Arntl*) deletion in the liver and its effect on REV-ERBa (*Nr1d1*) by RT-qPCR (n = 5-6, mean ± SEM). (**D**) Food intake patterns every hour over 24h in light:dark conditions (n = 3-4 mean ± SEM). (**E-F**) Total 24h food intake and percentage of 24h food intake in the light phase were calculated in BMAL1 deletion and control groups (n = 3-4, mean ± SEM). (**B-C & E**) Results were compared by one way ANOVA. (**D**) Results were compared with repeated measures ANOVA. (**F**) Results were analyzed using a one-way ANOVA test for trend. *p<0.05, **p<0.01, ***p<0.001, ****p<0.0001.

Although specific disruption of the hepatocyte molecular clock suggests that alterations in food intake are due to a signal emanating from the liver, surgical HVx disrupts both afferent and efferent signaling between the liver and brain. To address the role of afferent-specific signaling, we ablated vagal afferent neurons by injecting rgAAV-hSyn-Cre into the hepatic portal vein and a Cre-dependent caspase virus (or Cre-dependent mCherry virus as control) bilaterally into the nodose ganglia of HepDKO mice (**Fig. 5A**). Ablation of vagal afferent neurons by caspase prevented the aberrant food intake patterns of HepDKO mice (**Fig. 5B-D, Fig. S9A**), consistent with the hypothesis that afferent vagal signaling acts as mediator of the desynchronous signal from the liver with a genetically defective clock.

**Figure 5.**
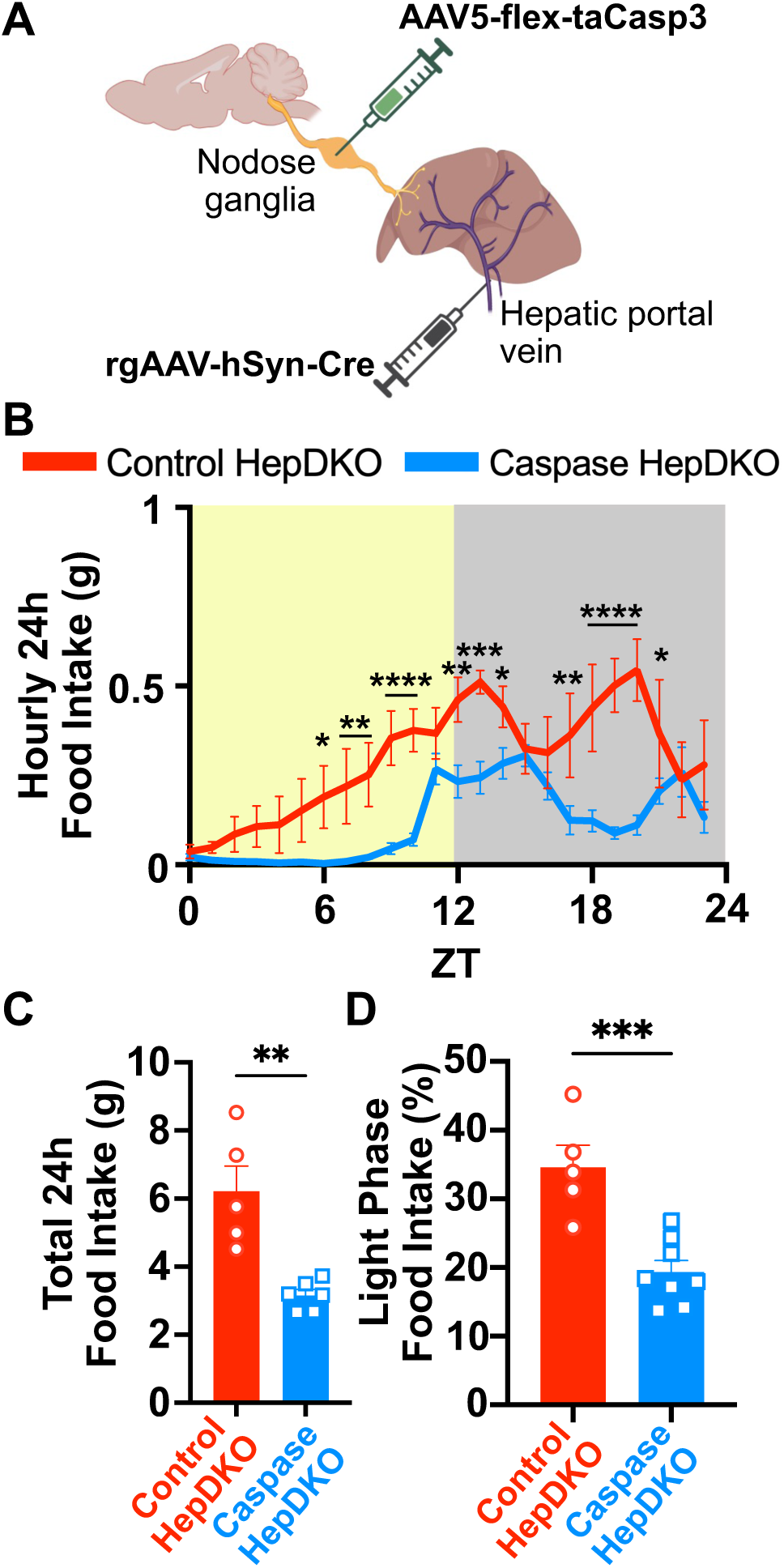
Disruption of vagal afferent signaling prevents aberrant food intake patterns from internal desynchrony induced by HepDKO. (**A**) HepDKO animals received hepatic portal vein injections of rgAAV-hSyn-Cre two weeks prior to receiving either AAV5-flex-mCherry (control) or AAV5-flex-taCasp3 (Caspase). (**B**) Food intake patterns every hour over 24hrs in control and Caspase groups (n = 5-8 mean ± SEM). (**C-D**) Total 24h food intake and percentage of 24h food intake in the light phase were calculated in control and Caspase groups (n = 5-8, mean ± SEM). (**B**) Results were compared by repeated measures. (**C-D**) Results were compared by Mann-Whitney U test. *p<0.05, **p<0.01, ***p<0.001 ****p<0.0001.

Intriguingly, high-fat diet (HFD) feeding and diet-induced obesity (DIO) have been shown to cause mice to consume a higher percentage of their daily food during the light/inactive phase (*30*, *31*). The mechanism by which this occurs is unknown, but the phenomenon is reminiscent of what we now observe with disruption of the liver clock and, interestingly, DIO is known to disrupt rhythmic gene expression and metabolism in peripheral tissues such as the liver (*30*, *32*). Therefore, we hypothesized that HVAN signaling may be involved in the altered food intake patterns observed in DIO animals. To test this, we performed Sham and HVx surgeries on naïve mice before placing on a HFD for 12 weeks. Sham animals developed robust disruption in food intake patterns and consumed nearly 50% of their daily food during the light phase (**Fig. 6A-C**) This HFD-induced shift in circadian food intake patterns was prevented by HVx (**Fig. 6A-C**). Moreover, the HVx animals gained less weight on HFD than their Sham counterparts (**Fig. 6D**).

**Figure 6.**
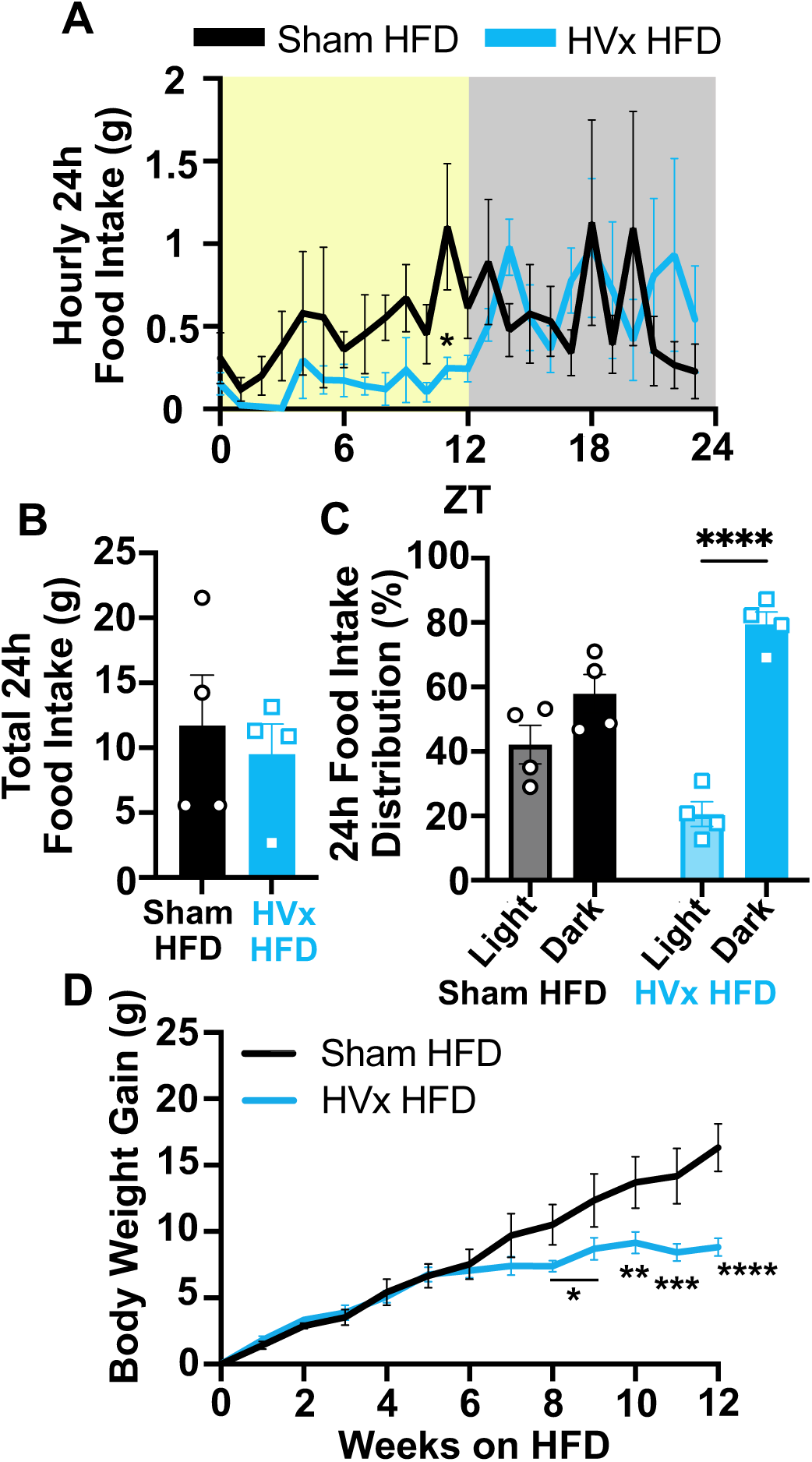
HVx protects against diet-induced disruptions in food intake patterns and body weight gain. (**A**) Food intake every hour over 24hrs measured in Sham and HVx animals maintained on a 60% HFD in light:dark conditions (n = 4, mean ± SEM). (**B-C**) Total 24h food intake and percentage of 24h food intake in the light and dark cycles were calculated in Sham and HVx HFD animals (n = 4, mean ± SEM). (**D**) Body weight gain over 12 weeks of HFD treatment in Sham and HVx animals (n = 4, mean ± SEM). (**B-C**) Results were compared by Mann-Whitney U test. (**A&D**) Results were compared by repeated measures ANOVA. *p<0.05, **p<0.01, ***p<0.001, ****p<0.0001.

Mammalian physiology has evolved to defend a set body weight to maintain fitness and survival in the face of varying environmental conditions. The 24-hour day is a prime example of environmental extremes against which an organism must maintain internal homeostasis. Indeed, light exposure synchronizes the central clock in the SCN whereas food acts as the Zeitgeber for the liver (*3*, *4*, *33*). Under normal conditions, the liver clock is synchronized to the SCN by physiological pattens of feeding that are driven by SCN-dependent behavioral and locomotor activity. By crippling hepatocyte clocks in two distinct ways (HepDKO and HepBKO) our data show that circadian disruption in the liver leads to desynchrony of a vagally-mediated signal that fed/fasting state does not match the light/dark environment. Feedback via the vagus, while normally homeostatic, signals internal desynchrony when the liver is disrupted by clock gene deletion, leading to maladaptive changes in food intake.

Food intake patterns in mice on HFD resemble those of mice with liver clock defects, and indeed we have found that HVx corrects daily feeding patterns and protects against DIO. This lends credence to and provides mechanistic insights into previous findings that HVAN blockade reduces body weight gain on HFD (*34*, *35*). Our findings have broad reaching implications for the obesity epidemic since healthy circadian rhythms are disrupted by poor diet and obesity (*10*). Consuming the majority of calories late in the day and/or night eating syndrome are correlated with obesity, poor metabolic outcomes and non-alcoholic fatty liver disease in human patients (*36–38*). Circadian disruption can also negatively impact metabolism such that individuals subjected to shift-work and jetlag are at an increased risk for developing obesity and metabolic disease (*7*).

Neuromodulation of the vagus nerve has been found to moderately improve body weight gain and metabolic outcomes in human patients (*39–41*). However, the results are variable, and the long-term efficacy is questionable due to the general modulation of function- and tissue-specific vagal fibers by current vagal neuromodulation techniques. In fact, work on gut vagal afferents indicates that activation, rather than inhibition, reduces food intake and improves metabolic outcomes (*42*). Here we show that the opposite is true for the hepatic vagus nerve and that blocking HVAN signaling prevents increased food intake in the face of internal circadian desynchrony. Therefore, stimulating or inhibiting the vagus nerve as a whole may confound the metabolic benefits of selectively modulating activity of specific vagal branches such as the hepatic branch, addressed in the present study. Our results indicate that developing technology to specifically inhibit hepatic vagal signaling may be a viable intervention for preventing the negative metabolic effects of diet-induced chrono-disruption between the brain and liver.

## Acknowledgements

The authors would like to thank M. Burrows, L. Pécout, R. Méndez-Hernández, T. Borner, T. Acavedo, M. Tackenberg, and all other members of the Lazar, Hayes, and de Lartigue labs for their invaluable help and discussion. We would also like to thank C. Holman and the Rodent Metabolic Phenotyping Core for their help with CLAMS experiments and the Penn Diabetes Research Center and Institute for Diabetes, Obesity, and Metabolism (DK19525).

## Funding

National Institutes of Health grant T32DK007314 (LNW)

National Institutes of Health grant F32DK128984 (LNW)

National Institutes of Health grant R01DK45586 (MAL)

National Institutes of Health grant R01DK105155 (MRH)

National Institutes of Health grant R01DK116004 (GL)

National Institutes of Health grant R01DK094871 (GL)

National Institutes of Health grant P30DK19525 (MAL)

JBP Foundation (MAL)

Cox Medical Institute (MAL)

## Author contributions

Conceptualization: LNW, GL, MRH, MAL

Methodology: LNW, AMA, CEG, GL, MRH, MAL

Investigation: LNW, LCM, MM, AMA, CEG, AJA, BMK, DMZ

Visualization: LNW, LCM, MM, AJA

Funding acquisition: LNW, GL, MRH, MAL

Project administration: LNW, GL, MRH, MAL

Supervision: GL, MRH, MAL

Writing – original draft: LNW, MAL

Writing – review & editing: LNW, LCM, MM, AMA, CEG, AJA, BMK, DMZ, GL, MRH, MAL

## Competing interests

The authors declare the following competing interests: MAL is on the advisory board and has received research funding unrelated to these studies from Pfizer, serves on the advisory board and is co-founder of Flare Therapeutics, and has consulted for Madrigal Pharmaceuticals. MRH receives additional research funding from Boehringer Ingelheim, Novo Nordisk, Pfizer, Gila Therapeutics, and Eli Lilly & Co. that was not used in support of these studies. LNW, LCM, MM, AMA, CEG, AJA, BMK, DMZ, and GL have no competing interests to declare.

## Data and materials availability

The GEO accession number for RNAseq data reported in this paper is GSE248462. All data will be made available upon request to LNW and MAL. MAL obtained the *Nr1d1* floxed mice under a material transfer agreement with Genoway. MAL obtained the *Nr1d2* floxed mice under a material transfer agreement with the Centre Europeen de Recherche en Biologie et Medecine. *Nr1d1^fl/fl^/Nr1d2^fl/fl^* mice are available by request to MAL.

## Materials and Methods

### Animals and diet

All animal work was approved by the University of Pennsylvania Perelman School of Medicine Institutional Animal Care and Use Committee (IACUC protocol number 804747 issued to Mitchell Lazar) in accordance with NIH guidelines. Male wildtype C57Bl/6J (Jackson Labs, #000664), *Nr1d1^fl/fl^/Nr1d2^fl/fl^*(lab stock), and *Arntl^fl/fl^* (Jackson Labs, #007668) mice were maintained on a C57Bl/6J background. G Power software was used to determine the minimum sample size necessary to achieve statistical significance at a=0.05 and b=0.2. Mice were bred and maintained at 20-22°C on a 12:12 hr light:dark cycle (lights on at 7am [Zeitgeber time 0, ZT0] and lights off at 7pm [ZT12]) with *ad libitum* access to food and water unless otherwise noted. The normal chow diet (NCD, LabDiet, 5010) consisted of 12.7% kcal fat, 28.7% kcal protein, and 58.5% kcal carbohydrates. The high-fat diet (HFD, Research Diets, D12492) consisted of 60% kcal fat from lard and soybean oil, 20% kcal protein, and 20% kcal carbohydrates. Surgical procedures and viral injections were performed on 6–8-week-old male littermates. To specifically delete REV-ERBs or BMAL1 in adult hepatocytes, adeno-associated virus serotype 8 (AAV8) encoding EGFP or Cre driven by the hepatocyte-specific TBG promoter (AAV8-TBG-EGFP for control and AAV8-TBG-Cre for knockout) were prepared by the UPenn Vector Core and were intravenously injected with 2×10^11^ genome copies (GC) per mouse. To determine genes disrupted by hepatocyte-specific REV-ERBs or BMAL1 deletion while minimizing batch effects, we collected tissues, extracted RNA, generated cDNA and/or RNAseq libraries for control and knockout samples at the same time. At each time point we alternated the sacrifice of control and knockout animal to avoid further bias. Micropunches of the Arc were pooled from three separate animals per group per time point for RNAseq. Four left nodose were pooled per group per time point for RNAseq.

### Common hepatic branch vagotomy

Surgery was performed aseptically following the IACUC guidelines for rodent survival surgery. Mice were anesthetized by inhalation of a continuous flow of 1-4% isoflurane. The pedal reflex was tested prior to surgery and at regular intervals throughout to ensure that the mice had reached an appropriate level of anesthesia. Analgesia was administered subcutaneously at the time of surgery (meloxicam 5mg/kg) and for three days post-op for pain management. Mice were placed on a sterile drape placed over a warming pad. Their fur was removed from the abdomen with depilatory cream (Nair) and cleaned with three changes of alternating antiseptic solution (Betaine) and ethanol. Sterile surgical equipment was used to create a 2-4cm midline laparotomy. The small intestine and colon were externalized and placed on sterile gauze moistened with sterile 0.9% saline. The common hepatic vagus nerve was visualized by gentle retraction of the liver and stomach. The nerve was isolated as it extends from the esophagus to the caudal aspect of the liver and sheared between two sterile cotton swabs to prevent regeneration. Sham animals had their vagus nerve visualized, but not tampered with. The liver, stomach, small intestine, and colon were then repositioned, and the incision was closed. Antibiotic ointment was applied to the site and the incision was secured shut by sterile wound clips. The animals’ recovery was monitored for ten days post-op and wound clips were removed fourteen days post-op. Animals were used for further experimentation only after wound clip removal.

### Hepatic portal vein injections

Surgery was performed aseptically following the IACUC guidelines for rodent survival surgery. Mice were anesthetized by inhalation of a continuous flow of 1-4% isoflurane. The pedal reflex was tested prior to surgery and at regular intervals throughout to ensure that the mice had reached an appropriate level of anesthesia. Analgesia was administered subcutaneously at the time of surgery (meloxicam 5mg/kg) and for three days post-op for pain management. Mice were placed on a sterile drape placed over a warming pad. Their fur was removed from the abdomen with depilatory cream (Nair) and cleaned with three changes of alternating antiseptic solution (Betaine) and ethanol. Sterile surgical equipment was used to create a 2-4cm midline laparotomy. The small intestine and colon were externalized and placed on sterile gauze moistened with sterile 0.9% saline. The hepatic portal vein (HPV) as it extends across the pancreas to the caudal aspect of the liver was visualized and isolated with forceps. 20μL of 1.3×10^13^ GC of rgAAV-hSyn-Cre (Addgene 105553) was mixed with 80μL of sterile 0.9% saline (100μL total injection volume) and slowly injected into the HPV to prevent flow into other tissues. Sterile gauze was applied over the site of injection as the needle was removed to reduce bleeding. The liver, stomach, small intestine, and colon were then repositioned, and the incision was closed. Antibiotic ointment was applied to the site and the incision was secured shut by sterile wound clips. The animals’ recovery was monitored for ten days post-op and wound clips were removed fourteen days post-op. Animals were used for further experimentation only after wound clip removal.

### Nodose injections

Surgery was performed aseptically following the IACUC guidelines for rodent survival surgery. Mice were anesthetized by inhalation of a continuous flow of 1-4% isoflurane. The pedal reflex was tested prior to surgery and at regular intervals throughout to ensure that the mice had reached an appropriate level of anesthesia. Analgesia was administered subcutaneously at the time of surgery (meloxicam 5mg/kg) and for three days post-op for pain management. Mice were placed on a sterile drape placed over a warming pad. Their fur was removed from the ventral aspect of the neck with depilatory cream (Nair) and cleaned with three changes of alternating antiseptic solution (Betaine) and ethanol. Sterile surgical equipment was used to create a 2cm midline incision and the underlying muscles, salivary glands, and lymph nodes were retracted. The nodose was identified by tracing the vagus nerve to the jugular foramen and additional muscles and connective tissues were retracted. A 30μm diameter tip glass micropipette was filled with either pAAV-hSyn-DIO-mCherry (Addgene 50459) or pAAV5-FLEX-taCasp3-TEVp (Addgene 45580). 0.5μL of the appropriate virus was injected into the nodose using a Picospritzer III injector (Parker Hannifin). The incision was closed with absorbable sutures and animals’ recovery was monitored for ten days. Animals were only used for further experimental procedures after fourteen days post-op.

### Metabolic rate and activity measurement

Metabolic rate and activity were measured by *in vivo* indirect calorimetry in Comprehensive Lab Animal Monitoring System (CLAMS, Columbus Instruments) cages. Animals were placed singly in CLAMS cages with bedding, food, and water that mimicked the home cage environment. Experimental data was collected after a two-day acclimation phase. Data were acquired every fifteen minutes by Oxymax software.

### Food intake measurement

Food intake was measured by Biological Data Acquisition (BioDAQ, Research Diets) cages. Animals were placed singly in BioDAQ cages with bedding, food, and water that mimicked the home cage environment. Experimental data was collected after a two-day acclimation phase. Data were acquired at each food hopper interaction and analyzed by BioDAQ Data Analysis software (**Table S1-5**).

### RNA extraction, cDNA synthesis, and RT-qPCR

For Arc RNA extraction, the area was microdissected from the hypothalamus and homogenized with 1mL of QIAzol Lysis Reagent (Qiagen, 79306) using Pellet-Pestel (Kimble, 749540). Liver was homogenized in 500uL of QIAzol Lysis Reagent using a TissueLyser system (Qiagen, 85300). After homogenization, total RNA from both tissues was purified and collected using RNeasy Mini Kits (Qiagen, 74004). RNA quality and quantity was determined by NanoDrop (Thermo Scientific, ND-ONE-W). cDNA was synthesized using High-Capacity cDNA Reverse Transcription kit (Applied Biosystems, 4368814). qPCR was run using Power SYBR Green PCR Master Mix (Applied Biosystems, 4368577) and QuantStudio 6 Flex Real-Time PCR system and software (Applied Biosystems, 4485691). All primers (**Table S6**) were validated, and RT-qPCR results were analyzed by standard curve and normalization to *Arbp* (liver) or *Tbp* (Arc and nodose).

### RNA sequencing

Micropunches of the Arc were pooled from three separate animals per group per time point for RNAseq. Four left nodose were pooled per group per time point for RNAseq. Total RNA was extracted (see RNA extraction, cDNA synthesis, and RTqPCR) from pooled Arc and pooled nodose samples. 500ng of total purified RNA was used for library preparation with the KAPA RNA HyperPrep Kit with RiboErase (HMR) for Illumina Platforms (KAPA Biosystems KK8561) according to the manufacturers protocol. Libraries were sent to Novogene, Inc. for sequencing on a NovaSeq X Plus.

### RNAseq data processing

RNA-sequencing reads were assessed for quality using FASTQC (0.11.7) and mapped to mouse genome (UCSC mm10) using STAR (V2.7.10b) with “--quantMode GeneCounts” to obtain gene-based counts. Normalized counts were obtained using varianceStabilizingTransformation function provided within DESeq2 within R 4.1.3. To identify the rhythmic genes for each tissue and condition, raw counts obtained from STAR were processed with dryR and JTK_CYCLE. KEGG pathway analysis was performed by uploading gene lists disrupted by HepDKO in the Arc or nodose to Enrichr (*43*). Significant pathways were selected for graphing and presentation on a -log10(p) scale.

### Data reproducibility and validation

Expression of molecular clock genes in Arc and nodose ganglia from control and HepDKO animals calculated from RNAseq data in Fig. 2C & 2G was validated by RT-qPCR in an independent cohort of mice (**Fig. S10A-B**).

### Statistical analysis

Statistical analysis and graphing were performed using GraphPad Prism 7 software (Dotmatics). All data were presented as mean ± standard error measurement (SEM) and statistical significance was set *a priori* as *p*<0.05. Statistical tests utilized to compare datasets are outlined in the legend of each figure.

---

**Figure S1.**
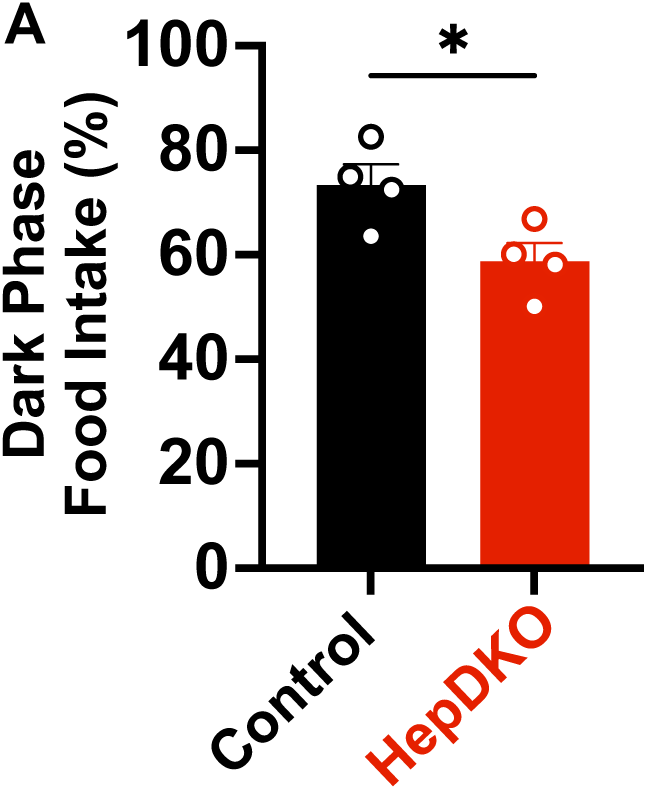
Control and HepDKO percent food intake in the dark phase. (**A**) Percentage of food intake during the dark phase in control and HepDKO animals (n = 4, mean ± SEM). Data were analyzed using Mann-Whitney U test. *p<0.05.

**Figure S2.**
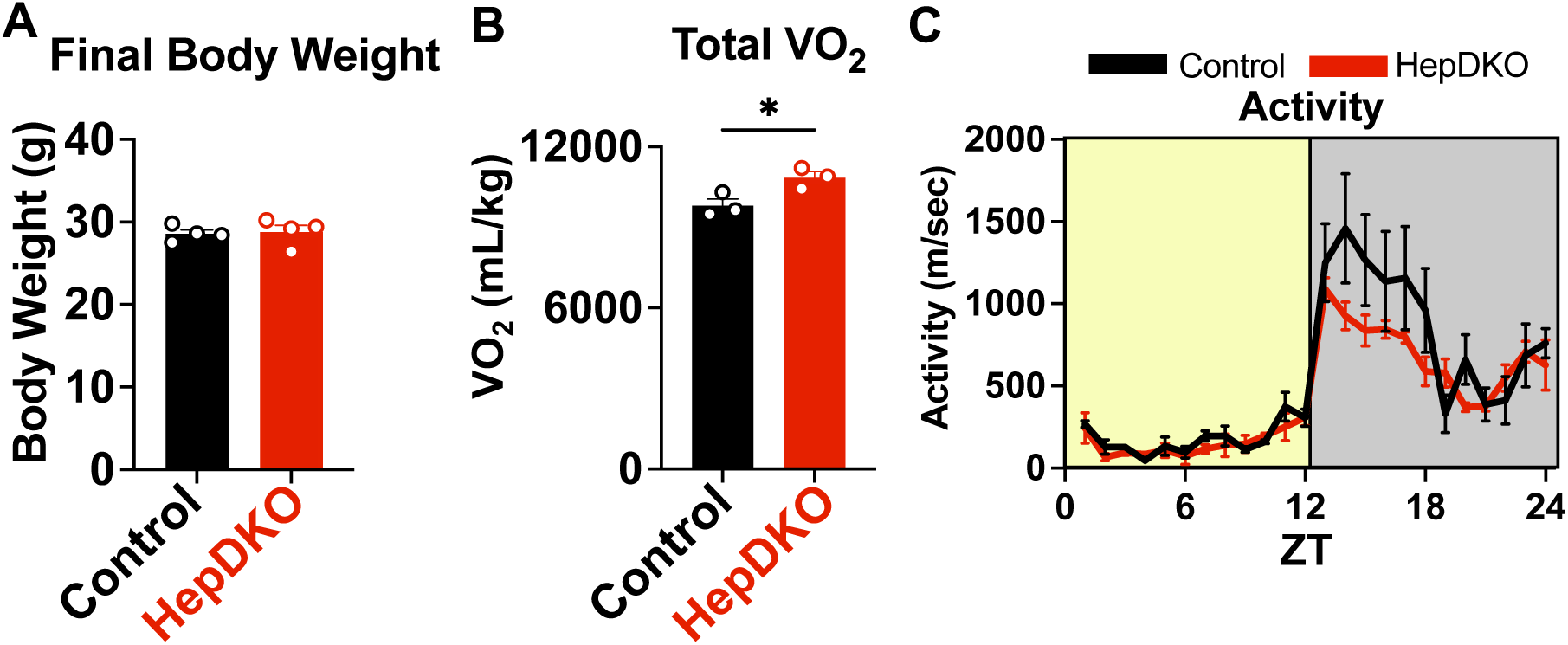
HepDKO phenotype. (**A**) Body weight at ZT10 sacrifice in control and HepDKO animals (n = 4, mean ± SEM). (**B**) Total VO_2_ over three consecutive days in CLAMS in control and HepDKO animals (n = 4, mean ± SEM). (**C**) Average of activity measured every hour over 24h in three consecutive days in control and HepDKO animals (n = 4, mean ± SEM). (**A-B**) Data were compared using Mann-Whitney U test. (**C**) Data were compared using repeated measured ANOVA. *p<0.05.

**Figure S3.**
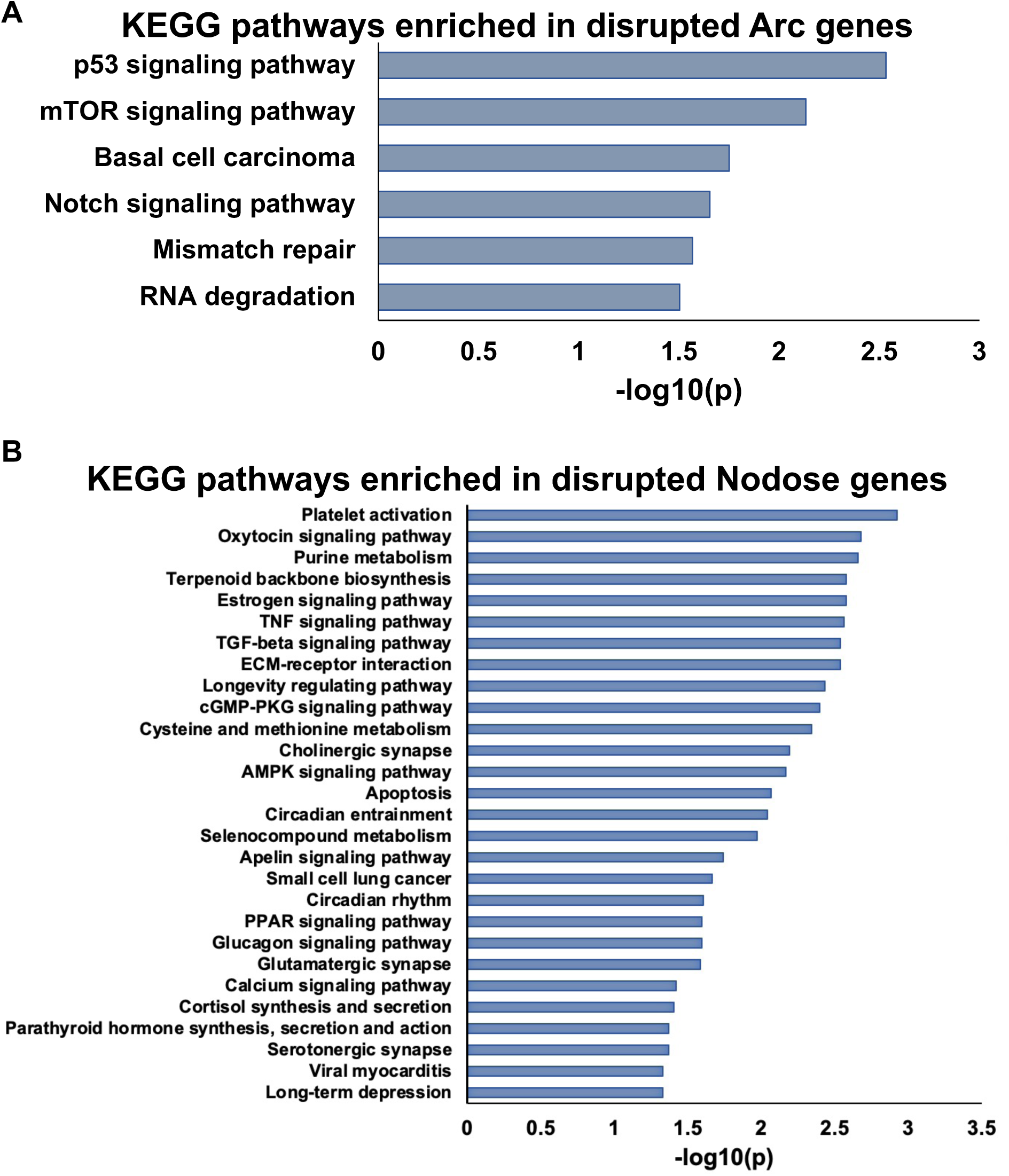
KEGG pathways enriched in genes disrupted by HepDKO in the Arc and nodose ganglia. (**A-B**) KEGG pathway analysis for pathways enriched in Arc and nodose genes disrupted by HepDKO. Disrupted genes in each tissue were uploaded to Enricher (*41*). Significantly enriched pathways (p<0.05) were selected for graphing as –log10(p).

**Figure S4.**
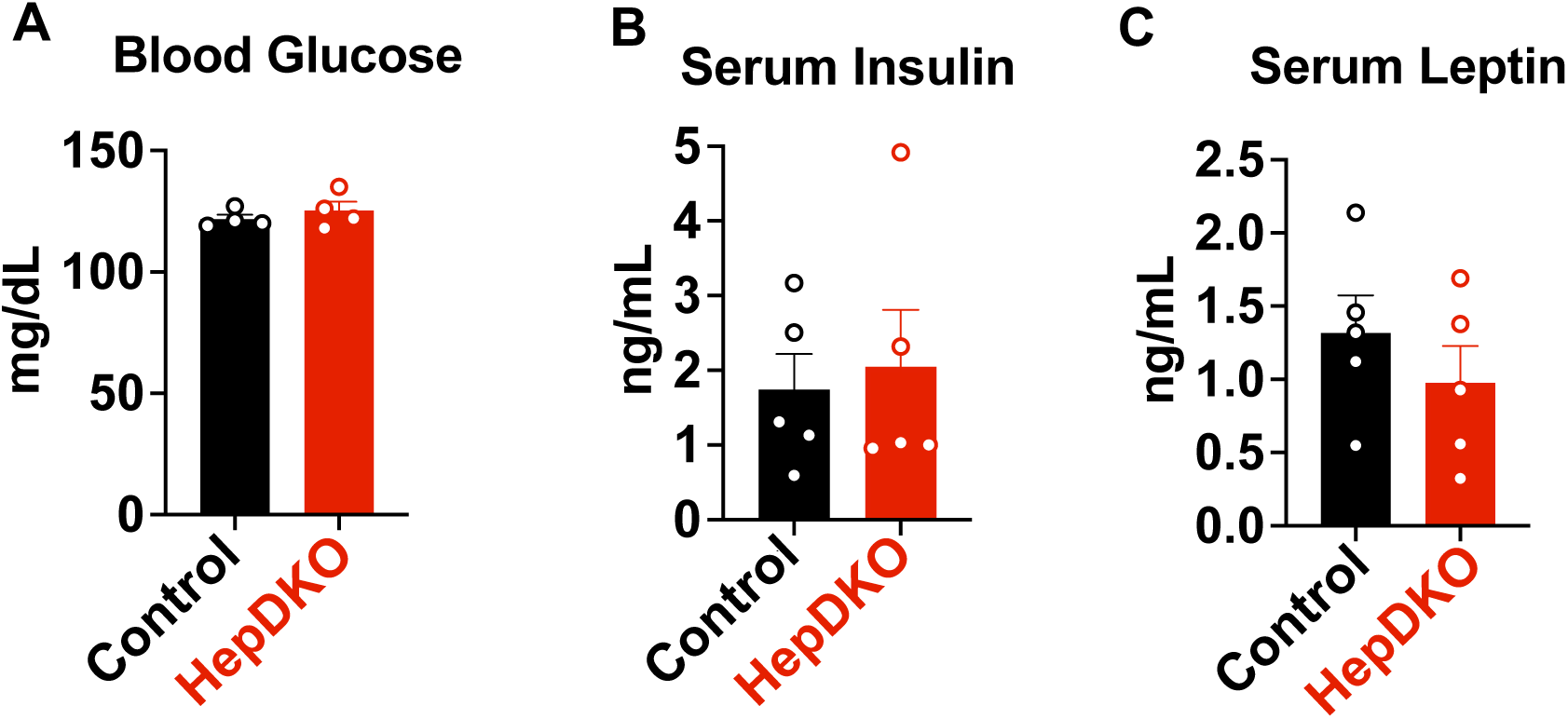
Circulating factors in control and HepDKO animals. (**A-C**) ZT10 blood glucose, serum insulin, and serum leptin in control and HepDKO animals (n = 4, mean ± SEM). Data were compared using Mann-Whitney U test.

**Figure S5.**
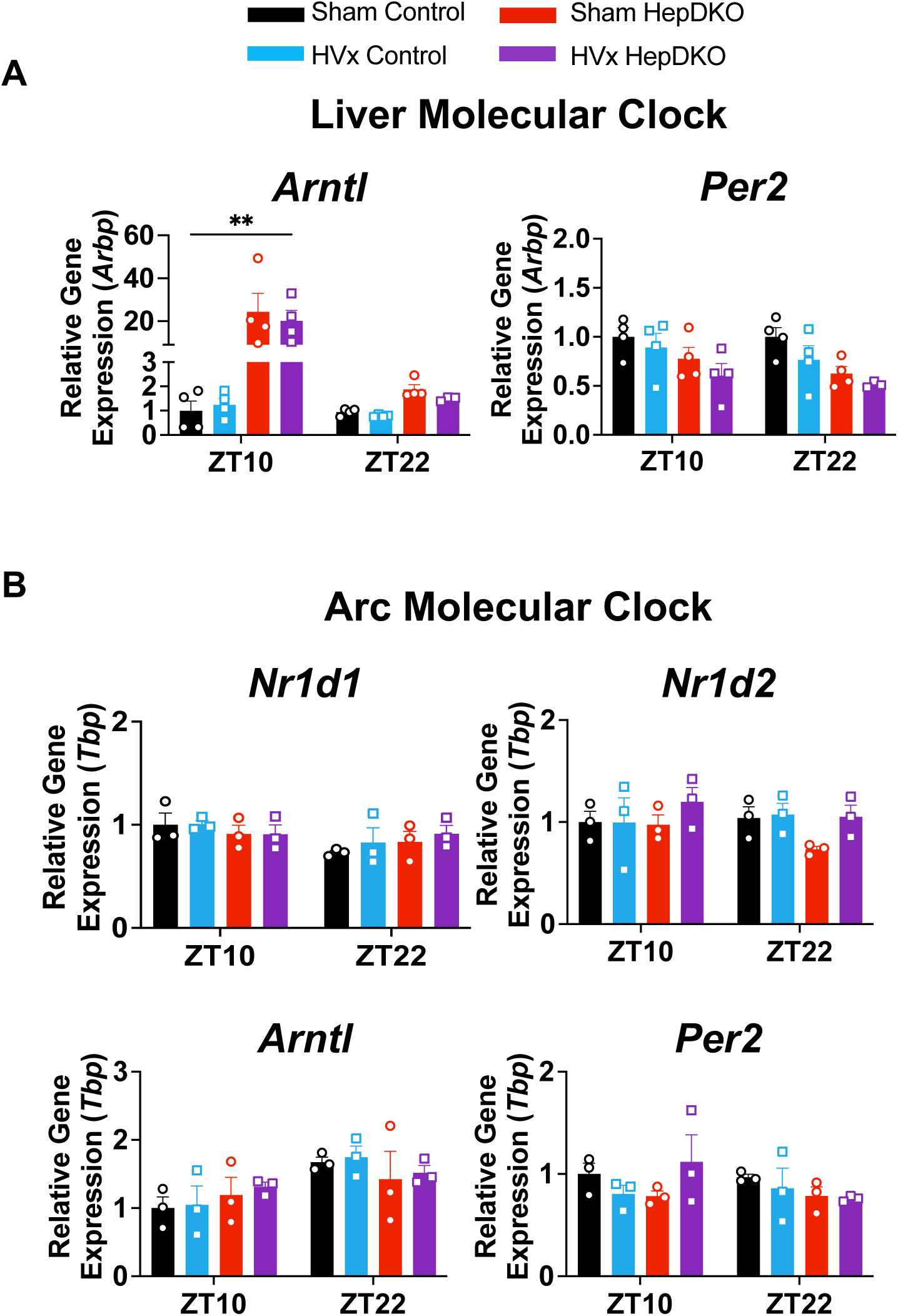
Sham and HVx HepDKO liver and Arc clocks. (**A**) RT-qPCR analysis of liver molecular clock genes in Sham control, HVx control, Sham HepDKO, and HVx HepDKO animals (n = 3-4, mean ± SEM). (**B**) RT-qPCR analysis of Arc molecular clock genes in Sham control, HVx control, Sham HepDKO, and HVx HepDKO animals (n = 3-4, mean ± SEM). (**A-B**) Data were analyzed using one-way ANOVA. **p<0.01.

**Figure S6.**
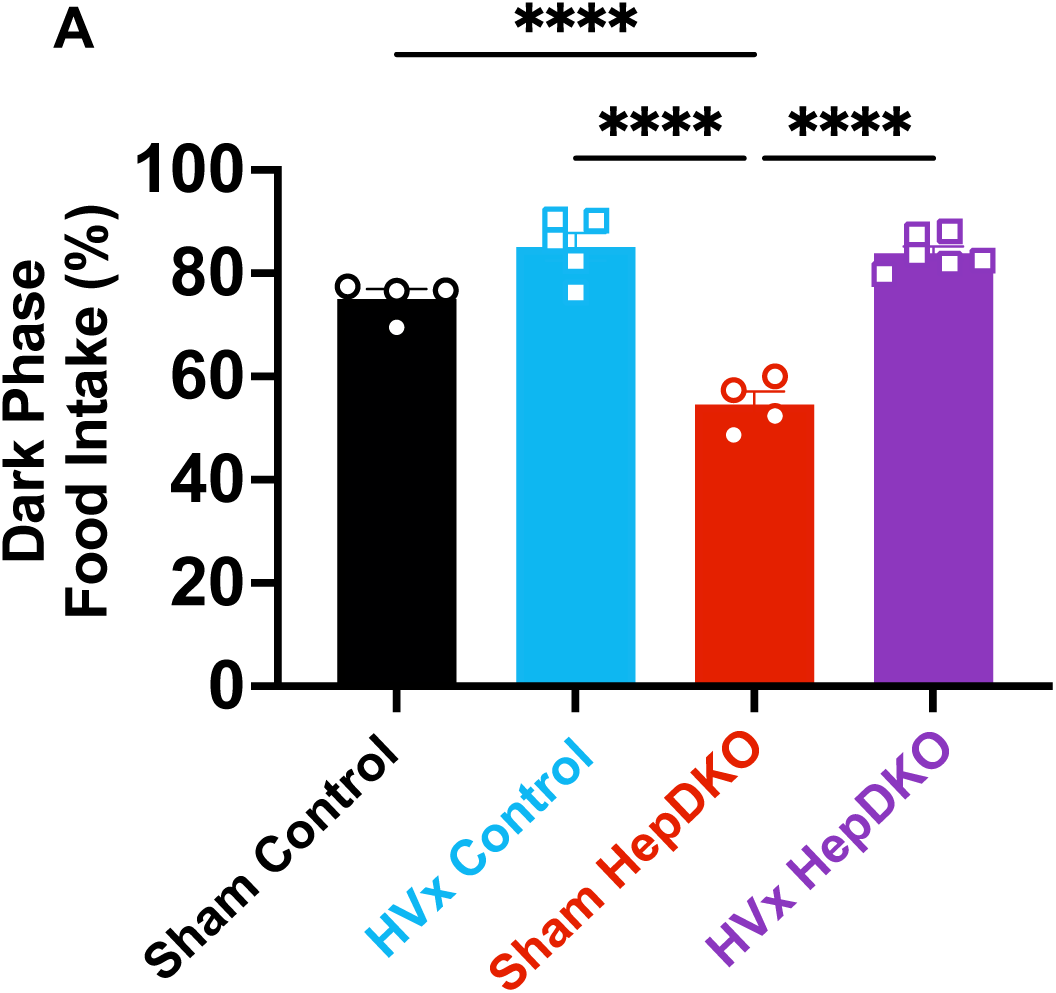
Sham and HVx HepDKO percent food intake in the dark phase. (**A**) Percentage of food intake during the dark phase in Sham control, HVx control, Sham HepDKO, and HVx HepDKO animals (n = 4-6, mean ± SEM). Data were analyzed using one-way ANOVA. ****p<0.0001.

**Figure S7.**
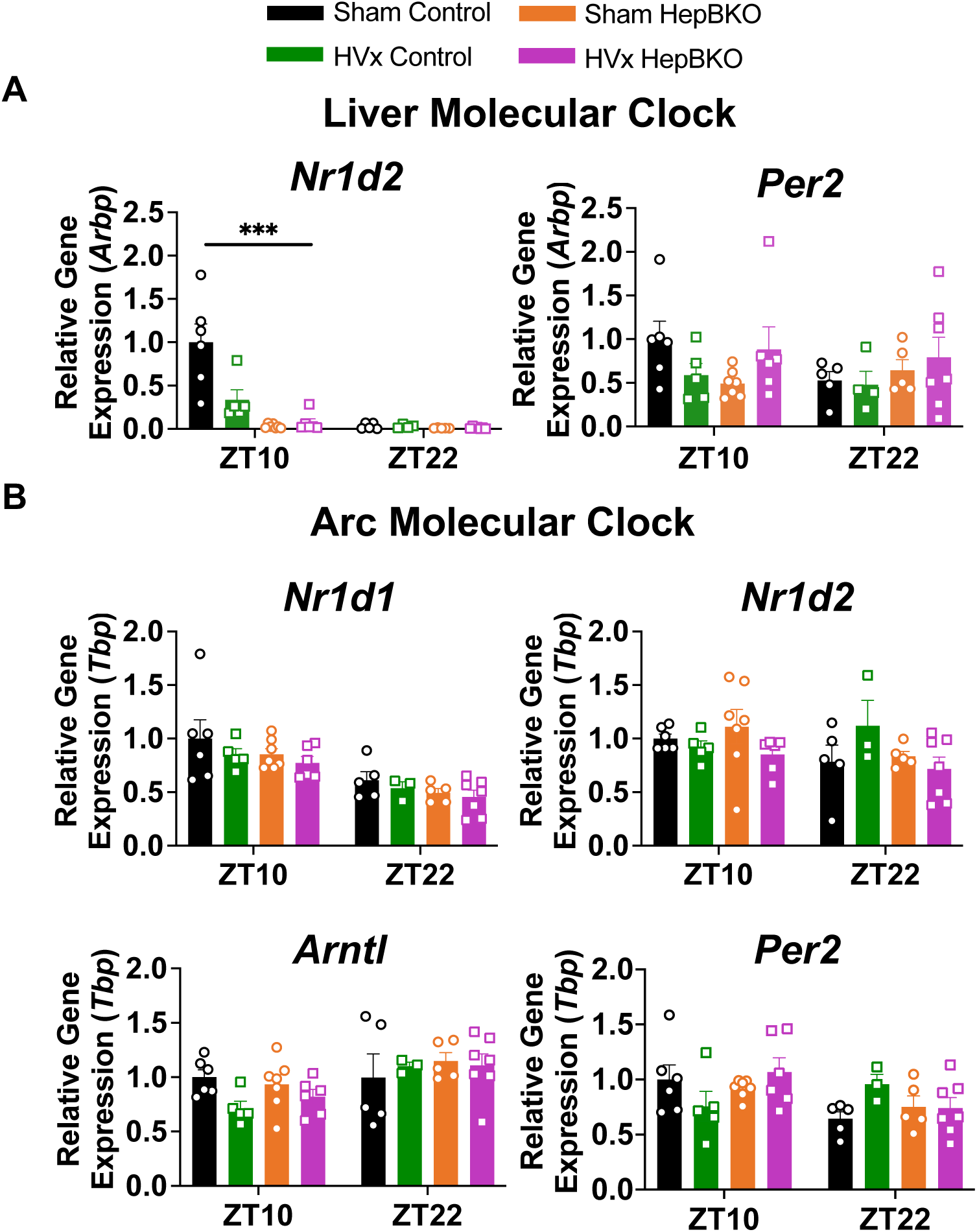
Sham and HVx HepBKO liver and Arc clocks. (**A**) RT-qPCR analysis of liver molecular clock genes in Sham control, HVx control, Sham HepBKO, and HVx HepBKO animals (n = 3-4, mean ± SEM). (**B**) RT-qPCR analysis of Arc molecular clock genes in Sham control, HVx control, Sham HepBKO, and HVx HepBKO animals (n = 3-4, mean ± SEM). (**A-B**) Data were analyzed using one-way ANOVA. ***p<0.001.

**Figure S8.**
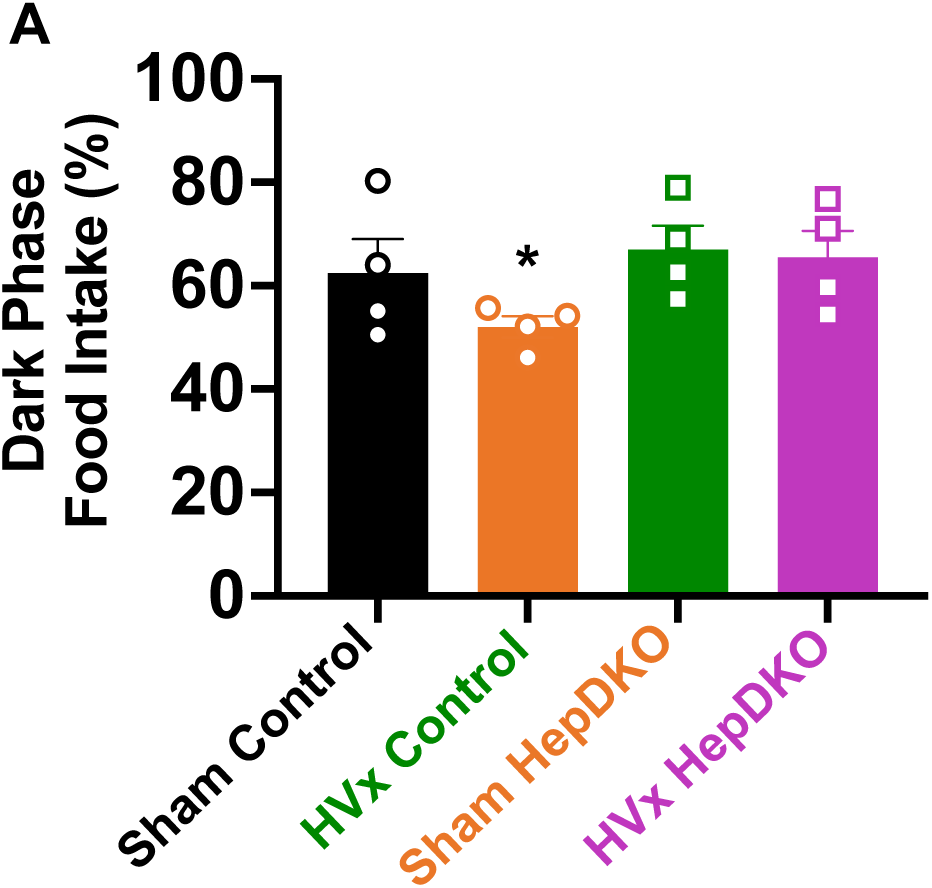
Sham and HVx HepBKO percent food intake in the dark phase. (**A**) Percentage of food intake during the dark phase in Sham Control, HVx Control, Sham HepBKO, and HVx HepBKO animals (n = 4, mean ± SEM). Data were analyzed using a one-way ANOVA test for trend. *p<0.05.

**Figure S9.**
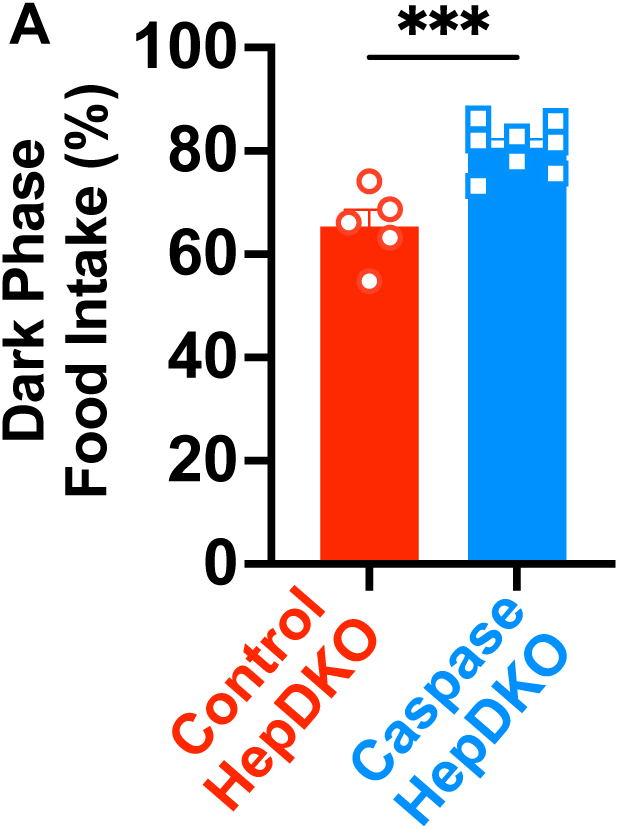
Control and Caspase HepDKO dark phase food intake. (**A**) Percentage of food intake during the dark phase in control and Caspase HepDKO animals (n = 3-4, mean ± SEM). Data were analyzed with Mann-Whitney U test. ****p<0.0001

**Figure S10.**
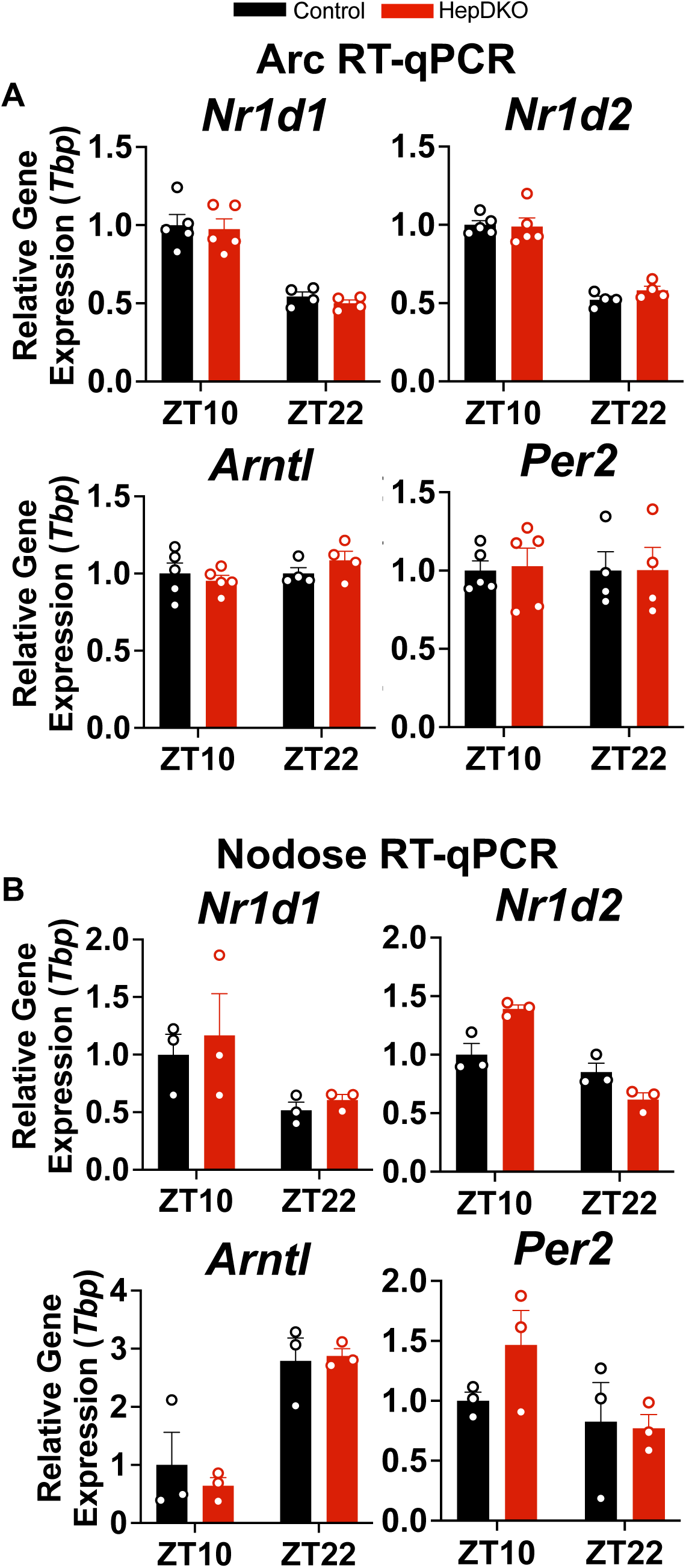
Data reproducibility and validation. (**A**) RT-qPCR analysis of Arc molecular clock genes in control and HepDKO animals validating RNAseq in Fig. 2C (n = 4-5, mean ± SEM). (**B**) RT-qPCR analysis of nodose molecular clock genes in control and HepDKO animals validating RNAseq in Fig. 2G (n = 3, mean ± SEM).

## References

1. C. Dibner, U. Schibler, U. Albrecht, The Mammalian Circadian Timing System: Organization and Coordination of Central and Peripheral Clocks. Annu. Rev. Physiol. 72, 517–549 (2010).

2. D. Guan, M. A. Lazar, Interconnections between circadian clocks and metabolism. J. Clin. Invest. 131 (2021), doi:10.1172/JCI148278.

3. D. Guan, Y. Xiong, T. M. Trinh, Y. Xiao, W. Hu, C. Jiang, P. Dierickx, C. Jang, J. D. Rabinowitz, M. A. Lazar, The hepatocyte clock and feeding control chronophysiology of multiple liver cell types Downloaded from. Science (80-. ). 369, 1388–1394 (2020).

4. K. A. Stokkan, S. Yamazaki, H. Tei, Y. Sakaki, M. Menaker, Entrainment of the circadian clock in the liver by feeding. Science (80-. ). 291, 490–493 (2001).

5. A. Atwood, R. DeConde, S. S. Wang, T. C. Mockler, J. S. M. Sabir, T. Ideker, S. A. Kay, Cell-autonomous circadian clock of hepatocytes drives rhythms in transcription and polyamine synthesis. Proc. Natl. Acad. Sci. U. S. A. 108, 18560– 18565 (2011).

6. R. van Ee, S. Van de Cruys, L. J. M. Schlangen, B. N. S. Vlaskamp, Circadian-Time Sickness: Time-of-Day Cue-Conflicts Directly Affect Health. Trends Neurosci. 39, 738–749 (2016).

7. J. P. Chaput, A. W. McHill, R. C. Cox, J. L. Broussard, C. Dutil, B. G. G. da Costa, H. Sampasa-Kanyinga, K. P. Wright, The role of insufficient sleep and circadian misalignment in obesity. Nat. Rev. Endocrinol. 2022 192. 19, 82–97 (2022).

8. A. Furlan, P. Petrus, Brain–body communication in metabolic control. Trends Endocrinol. Metab. 34, 813–822 (2023).

9. L. Small, L. S. Lundell, J. Iversen, J. T. Treebak, R. Barrè, J. R. Zierath Correspondence, A. M. Ehrlich, M. Dall, A. L. Basse, E. Dalbram, A. N. Hansen, J. R. Zierath, Seasonal light hours modulate peripheral clocks and energy metabolism in mice. Cell Metab. 35 (2023), doi:10.1016/j.cmet.2023.08.005.

10. L. N. Woodie, K. T. Oral, B. M. Krusen, M. A. Lazar, The Circadian Regulation of Nutrient Metabolism in Diet-Induced Obesity and Metabolic Disease. Nutr. 2022, Vol. 14, Page 3136. 14, 3136 (2022).

11. A. Bugge, D. Feng, L. J. Everett, E. R. Briggs, S. E. Mullican, F. Wang, J. Jager, M. A. Lazar, Rev-erbα and Rev-erbβ coordinately protect the circadian clock and normal metabolic function. Genes Dev. 26, 657–67 (2012).

12. N. Preitner, F. Damiola, Luis-Lopez-Molina, J. Zakany, D. Duboule, U. Albrecht, U. Schibler, The orphan nuclear receptor REV-ERBα controls circadian transcription within the positive limb of the mammalian circadian oscillator. Cell. 110, 251–260 (2002).

13. A. L. Hunter, C. E. Pelekanou, A. Adamson, P. Downton, N. J. Barron, T. Cornfield, T. M. Poolman, N. Humphreys, P. S. Cunningham, L. Hodson, A. S. I. Loudon, M. Iqbal, D. A. Bechtold, D. W. Ray, Nuclear receptor REVERBα is a state-dependent regulator of liver energy metabolism. Proc. Natl. Acad. Sci. U. S. A. 117, 25869– 25879 (2020).

14. M. Adlanmerini, B. M. Krusen, H. C. B. Nguyen, C. W. Teng, L. N. Woodie, M. C. Tackenberg, C. E. Geisler, J. Gaisinsky, L. C. Peed, B. J. Carpenter, M. R. Hayes, M. A. Lazar, REV-ERB nuclear receptors in the suprachiasmatic nucleus control circadian period and restrict diet-induced obesity. Sci. Adv. 7 (2021), doi:10.1126/SCIADV.ABH2007/SUPPL_FILE/SCIADV.ABH2007_TABLE_S2.ZIP.

15. C. M. Greco, K. B. Koronowski, J. G. Smith, J. Shi, P. Kunderfranco, R. Carriero, S. Chen, M. Samad, P. S. Welz, V. M. Zinna, T. Mortimer, S. K. Chun, K. Shimaji, T. Sato, P. Petrus, A. Kumar, M. Vaca-Dempere, O. Deryagian, C. Van, J. M. M. Kuhn, D. Lutter, M. M. Seldin, S. Masri, W. Li, P. Baldi, K. A. Dyar, P. Muñoz-Cánoves, S. A. Benitah, P. Sassone-Corsi, Integration of feeding behavior by the liver circadian clock reveals network dependency of metabolic rhythms. Sci. Adv. 7 (2021), doi:10.1126/SCIADV.ABI7828.

16. E. Challet, The circadian regulation of food intake. Nat. Rev. Endocrinol. 2019 157. 15, 393–405 (2019).

17. N. Sayar-Atasoy, I. Aklan, Y. Yavuz, C. Laule, H. Kim, J. Rysted, M. I. Alp, D. Davis, B. Yilmaz, D. Atasoy, AgRP neurons encode circadian feeding time. Nat. Neurosci. 2023, 1–14 (2023).

18. M. Quiñones, O. Al-Massadi, C. Folgueira, S. Bremser, R. Gallego, L. Torres-Leal, R. Haddad-Tóvolli, C. García-Caceres, R. Hernandez-Bautista, B. Y. H. Lam, D. Beiroa, E. Sanchez-Rebordelo, A. Senra, J. A. Malagon, P. Valerio, M. F. Fondevila, J. Fernø, M. M. Malagon, R. Contreras, P. Pfluger, J. C. Brüning, G. Yeo, M. Tschöp, C. Diéguez, M. López, M. Claret, P. Kloppenburg, G. Sabio, R. Nogueiras, p53 in AgRP neurons is required for protection against diet-induced obesity via JNK1. Nat. Commun. 9 (2018), doi:10.1038/S41467-018-05711-6.

19. D. Cota, K. Proulx, K. A. Blake Smith, S. C. Kozma, G. Thomas, S. C. Woods, R. J. Seeley, Hypothalamic mTOR signaling regulates food intake. Science (80-. ). 312, 927–930 (2006).

20. A. Jais, J. C. Brüning, Arcuate Nucleus-Dependent Regulation of Metabolism— Pathways to Obesity and Diabetes Mellitus. Endocr. Rev. 43, 314–328 (2022).

21. A. Kalsbeek, E. Fliers, A. N. Franke, R. M. Buijs, Functional Connections between the Suprachiasmatic Nucleus and the Thyroid Gland as Revealed by Lesioning and Viral Tracing Techniques in the Rat. Endocrinology. 141, 3832–3841 (2000).

22. C. X. Yi, S. E. la Fleur, E. Fliers, A. Kalsbeek, The role of the autonomic nervous liver innervation in the control of energy metabolism. Biochim. Biophys. Acta - Mol. Basis Dis. 1802, 416–431 (2010).

23. S. E. La Fleur, A. Kalsbeek, J. Wortel, R. M. Buijs, Polysynaptic neural pathways between the hypothalamus, including the suprachiasmatic nucleus, and the liver. Brain Res. 871, 50–56 (2000).

24. C. Cailotto, C. Van Heijningen, J. Van Der Vliet, G. Van Der Plasse, C. Habold, A. Kalsbeek, P. Pévet, R. M. Buijs, Daily Rhythms in Metabolic Liver Enzymes and Plasma Glucose Require a Balance in the Autonomic Output to the Liver (2008), doi:10.1210/en.2007-0816.

25. A. Pocal, T. K. T. Lam, R. Gutierrez-Juarez, S. Obici, G. J. Schwartz, J. Bryan, L. Aguilar-Bryan, L. Rossetti, Hypothalamic K(ATP) channels control hepatic glucose production. Nature. 434, 1026–1031 (2005).

26. T. K. T. Lam, A. Pocai, R. Gutierrez-Juarez, S. Obici, J. Bryan, L. Aguilar-Bryan, G. J. Schwartz, L. Rossetti, Hypothalamic sensing of circulating fatty acids is required for glucose homeostasis. Nat. Med. 2005 113. 11, 320–327 (2005).

27. V. Teckentrup, N. B. Kroemer, Mechanisms for survival: vagal control of goal-directed behavior. Trends Cogn. Sci. 0 (2023), doi:10.1016/J.TICS.2023.11.001.

28. H. R. Berthoud, V. L. Albaugh, W. L. Neuhuber, Gut-brain communication and obesity: understanding functions of the vagus nerve. J. Clin. Invest. 131 (2021), doi:10.1172/JCI143770.

29. H. R. Berthoud, W. L. Neuhuber, Functional and chemical anatomy of the afferent vagal system. Auton. Neurosci. 85, 1–17 (2000).

30. A. Kohsaka, A. D. Laposky, K. M. Ramsey, C. Estrada, C. Joshu, Y. Kobayashi, F. W. Turek, J. Bass, High-Fat Diet Disrupts Behavioral and Molecular Circadian Rhythms in Mice. Cell Metab. 6, 414–421 (2007).

31. A. Blancas-Velazquez, J. Mendoza, A. N. Garcia, S. E. la Fleur, Diet-induced obesity and circadian disruption of feeding behavior. Front. Neurosci. 11, 239991 (2017).

32. M. Hatori, C. Vollmers, A. Zarrinpar, L. DiTacchio, E. A. Bushong, S. Gill, M. Leblanc, A. Chaix, M. Joens, J. A. J. Fitzpatrick, M. H. Ellisman, S. Panda, Time-Restricted Feeding without Reducing Caloric Intake Prevents Metabolic Diseases in Mice Fed a High-Fat Diet. Cell Metab. 15, 848–860 (2012).

33. F. Damiola, N. Le Minh, N. Preitner, B. Kornmann, F. Fleury-Olela, U. Schibler, Restricted feeding uncouples circadian oscillators in peripheral tissues from the central pacemaker in the suprachiasmatic nucleus. Genes Dev. 14, 2950–61 (2000).

34. C. E. Geisler, S. Ghimire, C. Hepler, K. E. Miller, S. M. Bruggink, K. P. Kentch, M. R. Higgins, C. T. Banek, J. Yoshino, S. Klein, B. J. Renquist, Hepatocyte membrane potential regulates serum insulin and insulin sensitivity by altering hepatic GABA release. Cell Rep. 35 (2021), doi:10.1016/j.celrep.2021.109298.

35. C. E. Geisler, S. Ghimire, S. M. Bruggink, K. E. Miller, S. N. Weninger, J. M. Kronenfeld, J. Yoshino, S. Klein, F. A. Duca, B. J. Renquist, A critical role of hepatic GABA in the metabolic dysfunction and hyperphagia of obesity. Cell Rep. 35 (2021), doi:10.1016/j.celrep.2021.109301.

36. A. R. Gallant, J. Lundgren, V. Drapeau, The night-eating syndrome and obesity. Obes. Rev. 13, 528–536 (2012).

37. N. Perez-Diaz-Del-Campo, G. Castelnuovo, G. P. Caviglia, A. Armandi, C. Rosso, E. Bugianesi, Role of Circadian Clock on the Pathogenesis and Lifestyle Management in Non-Alcoholic Fatty Liver Disease. Nutrients. 14 (2022), doi:10.3390/nu14235053.

38. T. Marjot, J. W. Tomlinson, L. Hodson, D. W. Ray, Timing of energy intake and the therapeutic potential of intermittent fasting and time-restricted eating in NAFLD. Gut. 72, 1607–1619 (2023).

39. J. R. Fadel, L. P. Reagan, Stop signs in hippocampal insulin signaling: the role of insulin resistance in structural, functional and behavioral deficits. Curr. Opin. Behav. Sci. 9, 47–54 (2016).

40. F. V. Gouveia, E. Silk, B. Davidson, C. B. Pople, A. Abrahao, J. Hamilton, G. M. Ibrahim, D. J. Müller, P. Giacobbe, N. Lipsman, C. Hamani, A systematic review on neuromodulation therapies for reducing body weight in patients with obesity. Obes. Rev. 22, e13309 (2021).

41. H. R. Berthoud, W. L. Neuhuber, Vagal mechanisms as neuromodulatory targets for the treatment of metabolic disease. Ann. N. Y. Acad. Sci. 1454, 42–55 (2019).

42. G. de Lartigue, Role of the vagus nerve in the development and treatment of diet-induced obesity. J. Physiol. 594, 5791–5815 (2016).

43. Z. Xie, A. Bailey, M. V. Kuleshov, D. J. B. Clarke, J. E. Evangelista, S. L. Jenkins, A. Lachmann, M. L. Wojciechowicz, E. Kropiwnicki, K. M. Jagodnik, M. Jeon, A. Ma’ayan, Gene Set Knowledge Discovery with Enrichr. Curr. Protoc. 1, e90 (2021).

